# A naturally segregating polymorphism balancing semelparous reproduction versus reproductive diapause revealed via microfluidic assessment of starvation stress in *Caenorhabditis elegans*

**DOI:** 10.1101/863878

**Authors:** Heather Archer, Stephen Banse, Ben Blue, Patrick C. Phillips

**Affiliations:** Institute of Ecology and Evolution, University of Oregon, Eugene, Oregon, 97403

**Keywords:** *Caenorhabditis elegans*, GWAS, evolution, backcross, starvation, ecology

## Abstract

*Caenorhabditis elegans* typically feeds on rotting fruit and plant material in a fluctuating natural habitat, a boom-and-bust lifestyle. Moreover, stage specific developmental responses to low food concentration suggest that starvation-like conditions are a regular occurrence. In order to assess variation in the *C. elegans* starvation response under precisely controlled conditions and simultaneously phenotype a large number of individuals with high precision, we have developed a microfluidic device that, when combined with image scanning technology, allows for high-throughput assessment at a temporal resolution not previously feasible and applied this to a large mapping panel of fully sequenced intercross lines. Under these conditions worms exhibit a markedly reduced adult lifespan with strain-dependent variation in starvation resistance, ranging from <24 hours to ∼120 hours. Genome-wide mapping of the responses of more than 7,855 individuals identified four quantitative trait loci (QTL) of large effects. Three of these loci are associated with single genes (*ash-2, exc-6,* and *dpy-*28) and the fourth is a ∼26 KB region on Chromosome V encompassing several genes. Backcross with selection confirmed the effect of the Chromosome V locus. Segregating natural variation for starvation response in this species suggests that different isolates may use different strategies (facultative vivipary versus reproductive diapause) for dealing with extreme food deprivation.

## Introduction

In nature, overall physiology and behavior is governed by the energetic state of an organism, which in turn directly impacts how that organism invests its energetic resources when the availability of nutrients within the environment is uncertain. The behavioral response to environmental uncertainty typically manifests itself via trade-offs between investing energy in somatic maintenance and allocating resources toward reproduction. In particular, the nature of this balance may lead to delays or shifts in reproduction over the course of an individual’s life under conditions of reduced resource availability. For example, multiple bird species change how they invest in broods when resources are scarce, partitioning resources toward somatic development and delaying or foregoing reproduction for the season (Covas, Doutrelant, & du Plessis, 2004; Shaw & Levin, 2012; Mourocq et al., 2016). In stark contrast, some arachnid species address the challenge of limited resources via increased investment in reproduction to the point of facultative matriphagy (Stearns, 1989; Evans, Wallis, & Elgar, 1995; Kim, Roland, & Horel, 2000; Tizo-Pedroso & Del-Claro, 2005; Salomon, Aflalo, Coll, & Lubin, 2015). Other species have evolved to physically and temporally separate somatic growth and maintenance from reproduction, particularly in migratory birds and fish that leave their natal homes for more resource-rich environments before returning back to their breeding grounds and investing the acquired energy resources directly into reproduction (Stabell, 1984; Weber, Ens, & Houston, 1998; Bradbury et al., 2014; Moore et al., 2014). In its most extreme form, for instance in salmonids, the transition from somatic energy acquisition to reproductive investment is total, resulting in one-time, semelparous reproduction and death (Briggs, 1953; Kindsvater, Braun, Otto, & Reynolds, 2016).

Physiological effects of resource limitation can also more directly and immediately affect how individuals partition energy toward reproduction when nutrition is limited. For example, many mammalian species can delay implantation of fertilized eggs such that fetal development is postponed until nutrition is adequate, or for other species menstrual cycles become irregular and spontaneous abortion rates increase, and for unicellular organisms, nutritional state controls whether an individual develops to a reproductive state at all (Sandell, 1990; Bulik, et al., 1999; Trites & Donnelly, 2003; Kempes, Dutkiewicz, & Follows, 2011). The fact that such a broad diversity of species have evolved many independent mechanisms to partition resources in the face of nutrient limitation suggests that the trade-off between somatic and reproductive investment is common to most organisms. Currently missing, however, is a thorough understanding of the genetic and functional basis of the underlying systems involved in structuring that tradeoff.

As a model organism *Caenorhabditis elegans* has been widely used in laboratory settings for nearly 50 years (Brenner, 1974). In the past approximately 15 years, the natural ecology of *C. elegans* has begun to become more thoroughly understood (Félix & Braendle, 2010; Frézal & Félix, 2015). Breeding populations are typically found in decomposing vegetation in a continuum of settings ranging from human-affiliated (orchard and urban garden) to wild (forest and riverbank) (Félix and Braendle, 2010; Barriére and Félix, 2014). Collections of population samples from the same site over multiple seasons and years suggests a boom-and-bust pattern of population size variation. Genotypes have been observed to expand then shrink then expand, then appear to go extinct only to be followed by repopulation and inevitable subsequent disappearance (Félix and Braendle, 2010; Richaud, Zhang, Lee, Lee, & Félix, 2018). Given that *C. elegans* feeds on the bacteria associated with rotting plant material, which is itself subject to seasonal cycles in natural settings, it is likely that starvation conditions are a regular occurrence. Developmentally, *C. elegans* has unique stage specific responses to low food concentration which suggests that starvation-like conditions are naturally a regular occurrence in its environment and starvation a historically selective evolutionary pressure. If no food is present upon hatching, individuals arrest in the first larval stage (L1) only exiting that stage when sufficient food can be found (Johnson, Mitchell, Kline, Kemal, & Foy, 1984; Baugh, 2013; Roux, Langhans, Huynh, & Kenyon, 2016). If food is lacking early in the third (L3) or early in the fourth (L4) larval stage, development will halt in the respective stage (Schindler, Baugh, & Sherwood, 2014). If food runs low between the first and second larval stage (L2), *C. elegans* develops through the alternative dauer morph (Klass & Hirsh, 1976; Fielenbach & Antebi, 2008; Zhou, Pincus, & Slack, 2011; Roux, Langhans, Huynh, & Kenyon, 2016).

In addition to starvation induced larval arrest and induction of the dauer morph, a developmental response to starvation occurs in some individuals at the transition from the last larval stage (L4) to reproductive adult, termed adult reproductive diapause (ARD) (Angelo & Van Gilst, 2009; Seidel & Kimble, 2011; Burnaevskiy et al., 2018). If starvation occurs during this time, most individuals will continue to develop into reproductive adults and die via facultative matricide (aka “bagging”) because egg-laying is inhibited by starvation (Seidel & Kimble, 2011). However, in those worms that do not bag, the intestine and somatic gonad atrophy and the germline becomes reduced while other tissues continue to show signs of aging (Angelo and Van Gilst, 2009; Seidel and Kimble, 2011; Burnaevskiy et al., 2018). When food is reintroduced, these individuals exit ARD, growth resumes, germline stem cells repopulate, atrophy of the intestine and somatic gonad reverse, and animals become again capable of producing progeny, living a normal adult lifespan.

While there is extensive knowledge regarding *C. elegans* molecular and cellular biology along with its development and behavior, nearly all this information has come from a single, lab-domesticated genotype, largely ignoring the genetic diversity and natural variation present within the species and the related variety in phenotypes (Sterken, Snoek, Kammenga, & Andersen, 2015). Because the primary mode of reproduction is self-fertilization and the rate of outcrossing extremely low, populations sampled on the scale of <10 m often contain mostly individuals with identical or nearly identical genotypes (Barriére & Félix, 2005; Félix & Braendle, 2010). Even in samples taken from actively reproducing populations in which mixed genotypes were found, very few are heterozygous and, when examined repeatedly over a few years, no lasting effects of recombination was observed (Barriére & Félix, 2007; Rockman & Kruglyak, 2009; Andersen et al., 2012; Richaud, Zhang, Lee, Lee, & Félix, 2018). Recent simulations suggest these small-scale population dynamics are the result of demes founded by 3-10 individuals, suggesting the population structure is more appropriately depicted as ‘coexisting and competing homozygous clones’ (Richaud, Zhang, Lee, Lee, & Félix, 2018). Further, low levels of outcrossing lead to coadaptation of genomic loci. This results in selection acting against heterozygotes and/or recombinant genotypes and in fact outbreeding depression is frequent in *C. elegans* (Barriére & Félix, 2007; Dolgin, Charlesworth, Baird, & Cutter, 2007). So, while the use of a single genotype has facilitated many discoveries, the genetic diversity of *C. elegans* represents an interesting and largely untapped resource (Kammenga, Phillips, De Bono, & Doroszuk, 2008; Gaertner & Phillips, 2010; Frézal & Félix, 2015; Sterken, Snoek, Kammenga, & Andersen, 2015). Moreover, this natural population structure suggests that life history traits have the potential to be adapting via natural selection in parallel, sharing the same set of extrinsic factors while evolving under diverging internal constraints. Given the known differences in adult *C. elegans* responses to food availability, this is very likely the case in the context of starvation response.

Since the primary behavior of *C. elegans* when food becomes scarce is to leave the area, this typically results in individuals in a plate-based environment climbing up the walls of the plate where they desiccate and die, complicating phenotypic analysis of starvation response in these environments. Liquid-based assays can accommodate larger numbers without the loss to ‘walling’, but this leads to changes in behavior and physiology within the worms (Pierce-Shimomura et al., 2008). Moreover, both approaches are subject to the buildup of metabolic byproducts, which lead to the progressive alteration of the environment over time (Kaplan et al., 2011). This can be minimized by transferring individuals to fresh plates and/or conditions, but the physical handling and manipulation required introduces the possibility of stress and injury. The amount of labor required by these assays also limits the throughput and temporal resolution, such that most assays performed with larger samples are conducted with population cohorts and lack individual longitudinal information. Here, we develop a microfluidics approach that allows individual responses to be assayed at high temporal resolution using an automated scanner-based image acquisition system (based on Banse, Blue, Robinson, Jarrett, & Phillips, 2019). We are able to standardize the environment by using constant perfusion to flush metabolic byproducts and use an artificial dirt (pillared) structure (Lockery et al., 2008) so as to allow worms to display plate-like behavior and physiology.

We couple the scale, resolution, and throughput of microfluidics with whole-genome data for association mapping of adult starvation response in *C. elegans* using 72 recombinant inbred lines (RILs) from the *C. elegans* multiparental experimental evolution (CeMEE) panel (Noble et al., 2017). These lines were created by hybridization of 16 wild isolates, followed by 140 generations of mixed selfing and outcrossing then a further 50 generations selfing, all under standard laboratory conditions (Figure 1). The CeMEE panel captures 22% of the known polymorphisms segregating in wild populations and greater than 95% of the genome harbors nucleotide diversity. Moreover, intrachromosomal linkage decays to near background levels by 0.5 cM on average and interchromosomal disequilibrium is weak across chromosomes (Noble et al., 2017). In contrast, as mentioned above, natural populations have high linkage disequilibrium and the average SNP diversity is ∼0.3% (though in hypervariable regions it can reach >16%) (Cutter, 2006; Thompson et al., 2015). The low diversity and high linkage of natural populations can complicate standard GWAS approaches because correlated SNPs covering long genomic regions make it difficult to pinpoint loci relevant to the trait being studied and multiple correction testing becomes overly conservative (Graustein, Gaspar, Walters, & Palopoli, 2002; Rockman, Skrovanek, & Kruglyak, 2010; Andersen et al., 2012). Using the CeMEE panel lines, coupled with verification via a backcross with selection approach, we are able to examine the individual effects of natural genetic variants on adult starvation resistance at a finer resolution than possible in the presence of high linkage disequilibrium. We find that the genetic basis of adult lifespan under microfluidic starvation conditions is polygenic, with additive contributions from at least four loci, one which appears to be also pleiotropic for the ability to produce offspring after extended starvation.

**Figure 1:**
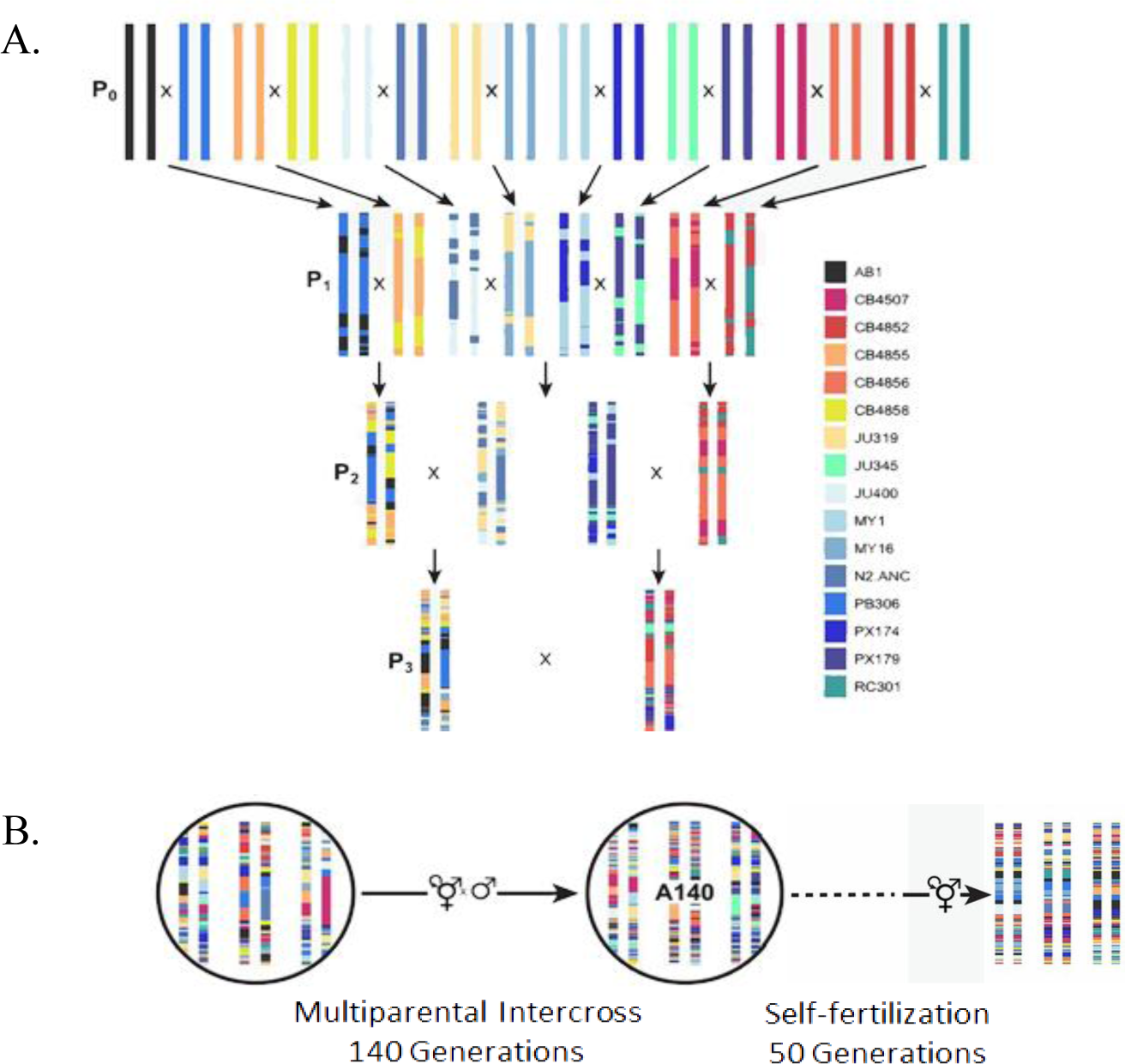
Derivation scheme for the C. elegans Multiparental Experimental Evolution (CeMEE) panel. The CeMEE panel captures 22% of the known polymorphisms segregating in wild populations and greater than 95% of the genome harbors nucleotide diversity. Intrachromosomal linkage decays to near background levels by 0.5 cM on average and interchromosomal disequilibrium is weak (r^2^ 0.99, 0.95 quantiles = 0.538, 0.051 within chromosomes vs. 0.037, and 0.022) across chromosomes. Adapted from Noble et al., 2017.

## Materials And Methods

### Mapping Lines

All *C. elegans* strains were maintained with standard culturing conditions on NGM lite agar with *E. coli* OP50 as a food source and maintained at 20°C (Brenner, 1964). The strains used for genetic mapping are recombinant inbred lines (RILs) from the *C. elegans* multiparental experimental evolution (CeMEE) panel (Noble et al., 2017). The strains used here are listed in Supplemental Table I.

### Phenotyping of starvation response using microfluidics

The Starvation Arena (Figure 2) was designed with Vectorworks Fundamentals (Vectorworks, Inc.). Using standard soft lithography methods (Whitesides, Ostuni, Takayama, Jiang, & Ingber, 2001), single layer devices were fabricated with polydimethylsiloxane (PDMS) and bonded to a glass microscopy slide by air plasma exposure (see Banse, Blue, Robinson, Jarrett, & Phillips, 2019). A biopsy punch was used to open inlet and outlet channels. Embryos were harvested via bleaching and immediately placed onto OP50 seeded plates at a density of ∼5000 individuals/100 mm petrie plate. When populations reached the L3 larval stage, additional OP50 was added in excess so individuals could feed ad libitum. Populations were monitored until ∼3 early adults were visible. At this time the majority of individuals in the population were in either very late L4 larval stage or extremely early adulthood. Hermaphrodites were immediately picked into a prepared microfluidic chip (see section below) at a density of ∼1 individual per arena (50 arenas per chip), and image acquisition initiated. All starvation experiments were performed in S Basal lacking cholesterol (5.85g NaCl, 1g K_2_HPO_4_, 6g KH_2_PO_4_, H_2_O to 1 liter) sterilized by autoclaving.

**Figure 2:**
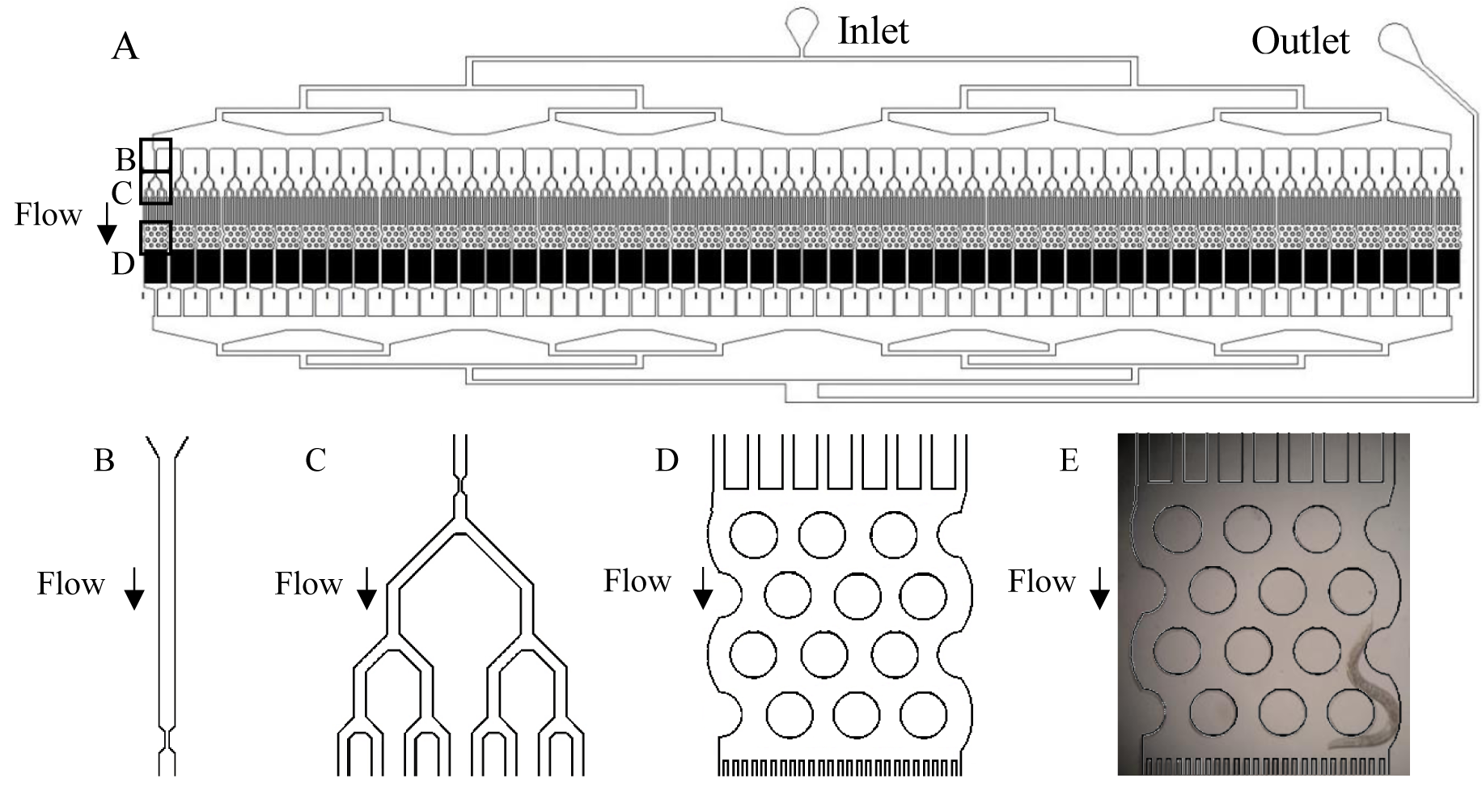
Microfluidic chip design. A) The device is designed to have a single array of 50 pillared arenas. Flow through the device enters at the inlet hole, perfuses through the device, and exits at the outlet hole. B) Movement of worms into the arenas is facilitated by a 0.52 mm upstream holding channel (1 per arena) with a 0.015 mm constriction point. This enables pre-loading of 50 animals followed by a brief increase in pressure sufficient to move the animals into the arenas. C) Each arena has a bifurcated section preceding the pillared portion to keep animals in the arenas. D) The arena is ∼1.2 mm x ∼0.9 mm with twelve 0.2 mm diameter pillars spaced 0.1 mm apart. E) An individual arena housing an adult *C. elegans* hermaphrodite.

### Experimental Protocol (Microfluidic Device Setup)

Four hours before use, microfluidic starvation arenas (aka chips) were first filled with an 1% w/v solution of Pluronic F-127 dissolved in S-basal made w/o cholesterol then allowed to sit for ∼40 minutes. This procedure lowers adhesion to the PDMS surface (Wu, 2009). Care was taken to ensure that no air was allowed into the chips from this point forward. The chips were then flushed with S-basal lacking cholesterol and ∼50 worms picked into the inlet port of each device. Tubing was connected to the chips and additional S-basal flushed through under manually controlled pressure until all worms were aligned properly in the array formation at the constriction point (Figure 2). Then, pressure was manually increased to a level sufficient to cause the expansion of the constriction point such that worms pass through and into the pillared portion of the arena. Lifespan assays of *C. elegans* using the same constriction point in a similar device but done in the presence of food result in a normal lifespan suggesting the passage of worms through the constriction point in this manner does not result in injury (Banse, Blue, Robinson, Jarrett, & Phillips, 2019).

Once worms had moved into the arenas, chips were placed on flatbed document scanners (Epson V700, model B11B178011) and connected to a mechanically pressurized fluid system with inflow air pressure set at 3 psi (Figure 3). To ensure that pressure stayed consistent throughout the course of a given experiment, an intermediate air tank was used as a buffer. In this system the pressure source coming in to the intermediate tank was ∼80 psi and subject to potential transient increases or decreases as the initiating compressor turned on and off. In order to buffer against these potential fluctuations and reduce the pressure to an appropriate level, a regulator was placed on the outflow side of the intermediate air tank such that the flow of fluid into the chips was ∼ 3 psi (Figure 3).

**Figure 3:**
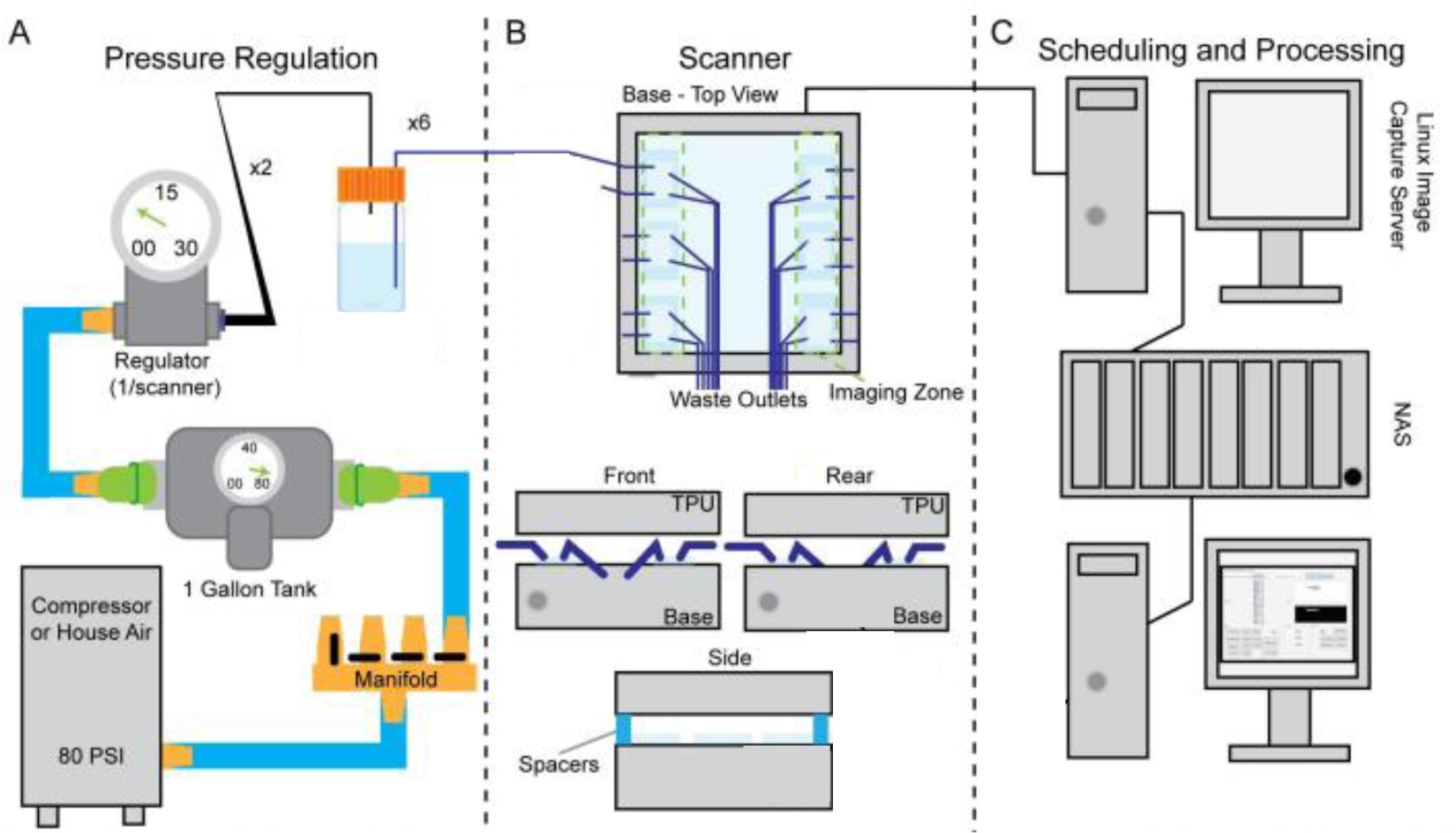
Pressure regulation and image capture schematic. A) Pressurized air was split through a manifold into a 1-gallon cushion tank. Air exiting the cushion tank was regulated with a microregulator set to 3-4 psi. This air was then passed through bifurcated tubing (Rain Bird ¼” blank tubing and ¼” barbed couplers) and into four 1-liter bottles. Each bottle lid had four 1.5 mm OD stainless steel stems passing through. One stem connected the pressurized air and the other three were connected internally to ∼50 mm of tubing extending to the bottom of the bottle, and externally to 1.5 mm ID tubing extending to the inlet ports of the microfluidic chips. B) Microfluidic chips bonded to 50 mm x 75 mm glass slides were arranged on scanner with 3 slides per side (2 chips per slide), ∼15 mm from the scanner edge. C) Image capture was done at timed intervals using a Linux version of the *C. elegans* Lifespan Machine software (Stroustrup et al., 2013) with images stored on a local network storage device. Image processing was performed using ImageJ 1.50i (Schneider, Rasband, & Eliceiri, 2012). Adapted from Banse, Blue, Robinson, Jarrett, & Phillips, 2019.

The air-line exiting the buffer tank was bifurcated into 4 tubing lines, each pressurizing a sealed 1 L bottle of S-basal made without cholesterol as previously described. The bottle caps had been modified with 4 ports: 1 terminal inlet port for air tubing and 3 exit ports for liquid supply lines. Each liquid supply line extended from the basal surface of the 1 L bottle to an inlet port of a microfluidic chip placed on a scanner (Figure 3). Once all chips were connected and fluid flow visually verified by monitoring the outflow through the arena device exit ports, image acquisition was initiated. Fluid flow was continuous throughout the course of any given experiment.

### Image Analysis and Survival Estimation

Scans of an individual microfluidic chip were initiated every 4 minutes in a sequential manner resulting in a rate of 1 image every 48 minutes for each of 12 chips per scanner. Images were collected for a period of 5 days using *C. elegans* Lifespan Machinery software (Stroustrup, et al. 2013; for additional technical details see Banse, Blue, Robinson, Jarrett, & Phillips, 2019).

Image processing was performed using ImageJ 1.50i (Schneider, Rasband, & Eliceiri, 2012). TIFF images were downloaded as a stack (one stack per chip) with brightness and contrast adjusted as necessary. Image stacks were manually scanned and worms examined individually. An individual was determined dead when movement was no longer visible for a minimum period of 4 frames (3 hours and 12 minutes). Individuals still alive at the conclusion of the experiment were censored and included in this manner. Analysis of Kaplan-Meier survivorship curves with log rank test for statistical differences was performed using JMP Statistical Software (SAS Institute Inc., JMP Pro 13, 2017). Possible differences in the shape of survivorship curves were assessed using the fit to a Gompertz mortality model using the R package flexsurv (Jackson, 2016).

### Genome Wide Association Test

The CeMEE strains used here contained 352,665 SNPs spread across 72 recombinant inbred lines (Noble et al., 2017, Supplemental Table 1). Using Plink v.1.9, SNPs with more than 10% missing calls across the lines were filtered from the analysis (www.cog-genomics.org/plink/1.9/; Chang et al., 2015). Further, individual lines were required to have genotypes at ≥ 95% of sites to be retained. GWAS analysis of survival time in the absence of food was conducted on the trimmed data using an additive linear regression in Plink v.1.9, using the following options: --assoc qt-means --adjust gc. Plink binary files were converted to the rrBLUP format. GWA mapping was performed in rrBLUP using the above genotype data and the quantitative trait of time-until-death for each individual in the microfluidic starvation environment using the standard GWAS function (GWAS(phenotype, genotype, min.MAF = 0.05, plot = TRUE) (Endelman, 2011). A kinship matrix was calculated by default with the A.mat function because the argument ‘K’ was not passed. SNPs with a minor allele frequency of less than 5% were excluded. After applying these filters, a total of 248,374 SNPs and 7,855 individuals remained.

### Backcross with Selection

In order to independently test the mapping of potential QTL influencing starvation response, a backcross with selection approach was implemented using two CeMEE lines that were found to display highly divergent starvation response phenotypes: A6140L110 as the ‘short-lived’ parent and GA450L40 as the ‘long-lived’ parent. Since males arise from non-disjunction of the X chromosome in meiosis and occur at a very low frequency in *C. elegans*, male-enriched lines were created by passaging an equal number of males to hermaphrodites (∼20 each) every generation until a sufficient number of males was consistently present (∼5 generations and ∼20% male frequency). After this, 25 day 1 adult hermaphrodites were selected from each parent line (n = 50), picked to an isolated plate (1 line per plate), and allowed to self-fertilize for 24 hours in order to deplete self-sperm. Once this period had passed, hermaphrodites were transferred to a new plate and an equal number of males from the opposing parent line were added. After ∼48 hours, eggs (F1 offspring) were harvested through treatment with sodium-hypochlorite (Stiernagle, 2006) and immediately placed on OP50 seeded plates. To allow for the segregation of any potential recessive alleles, all odd-numbered generations were allowed to develop without interference and self-fertilize, with eggs harvested as previously described.

Even-numbered generations, assumed to be ¼ homozygous for loci of interest, were allowed to develop to late L4/early adulthood and picked into microfluidic chips in the same manner as previously described under the Phenotyping of Starvation Response Using Microfluidics section but with one critical difference: pressure was not increased to a level sufficient to open the constriction point and move individuals into the arenas. Worms were kept upstream of the constriction point so surviving individuals could be recovered as follows: 1) chips were placed on scanners with image acquisition initiated as previously described, 2) images were monitored such that chips could be removed from scanners when approximately 20% of individuals remained alive or 24 hours had passed, whichever came first, 3) after removal, tubing was connected in reverse so the direction of fluid flow was reversed and the remaining individuals could be manually flushed out through the inlet ports and picked onto OP50 seeded plates.

Survivors were allowed to recover/self-fertilize on OP50 seeded plates for a period of 24 hours. After this, they were picked to new plates and males from the ‘short-lived’ parent line were added in a 1:1 ratio. Eggs were harvested via bleaching after ∼48 hours and the resulting odd-numbered generation was handled as previously described. This cycle was repeated up to and through the F11 generation.

Following thirteen generations of the above procedure (five generations of actual backcrossing), hermaphrodites were subjected to selection one final time and twenty survivors picked to individual plates as founders for near-isogenic lines (NILs, n = 20). Each of these lines was maintained in isolation and self-fertilized for 10 generations before being phenotyped again.

### Genomic Sequencing of NILs

Nematodes were maintained under standard lab conditions as described (Brenner, 1974). For preparation of genomic DNA, ∼10 - 20,000 synchronized L1 staged worms in M9 media were processed with the Zymo genomic DNA kit following Proteinase K digestion for 3-4 hours at 55°C. Genomic DNA concentration was quantified with a Qubit fluorimetric reader using the HS kit. Genomic DNA was processed for sequencing with the Nextera DNA Library kit per manufacturer instructions. Individual samples were multiplexed, combined in equal molar ratios and sequenced on a Hi-Seq 4000 with 100bp of single end reads. (University of Oregon Sequencing Facility, Eugene, OR). Variant calls were generated with GATK and line comparisons done with bcftools-isec (see supplemental files for detailed scripts).

### Data Availability

Sequence data for CeMEE panel lines are available from the National Center for Biotechnology Information Sequence Read Archive under BioProject PRJNA381203. Sequence data for NIL lines are under BioProject PRJNA589469. Raw phenotype data, supplemental figures/tables, data processing scripts, and NIL variant call files are available as supplemental material archived in FigShare at DOI 10.6084/m9.figshare.10314371.

## Results

### Variation in Adult Lifespan of CeMEE Panel Lines under Microfluidic Starvation Conditions

We assayed 72 CeMEE panel lines for starvation response using our novel microfluidic approach (Figure 4). Each CeMEE line represents a unique combination of the genetic variants segregating within the derived ancestral population (Figure 1). Median survival times vary by up to six-fold (range from 14.1 hours to 86.5 hours, Supplemental Table I). Partitioning variation within and between lines using analysis of variation of the survival time of each of the 7,855 individuals in the experiment reveals that genotype accounts for 35% of the total variation observed in the experiment (Restricted Maximum Likelihood, genotype and replicate nested within genotype as random variables, upper 95% CI = 265.52, lower 95% CI = 116.58, SE = 38.00, p < 0.0001). Overall, there appears to be substantial variation in both median survival time and in the shape of the mortality trajectories of each line.

**Figure 4:**
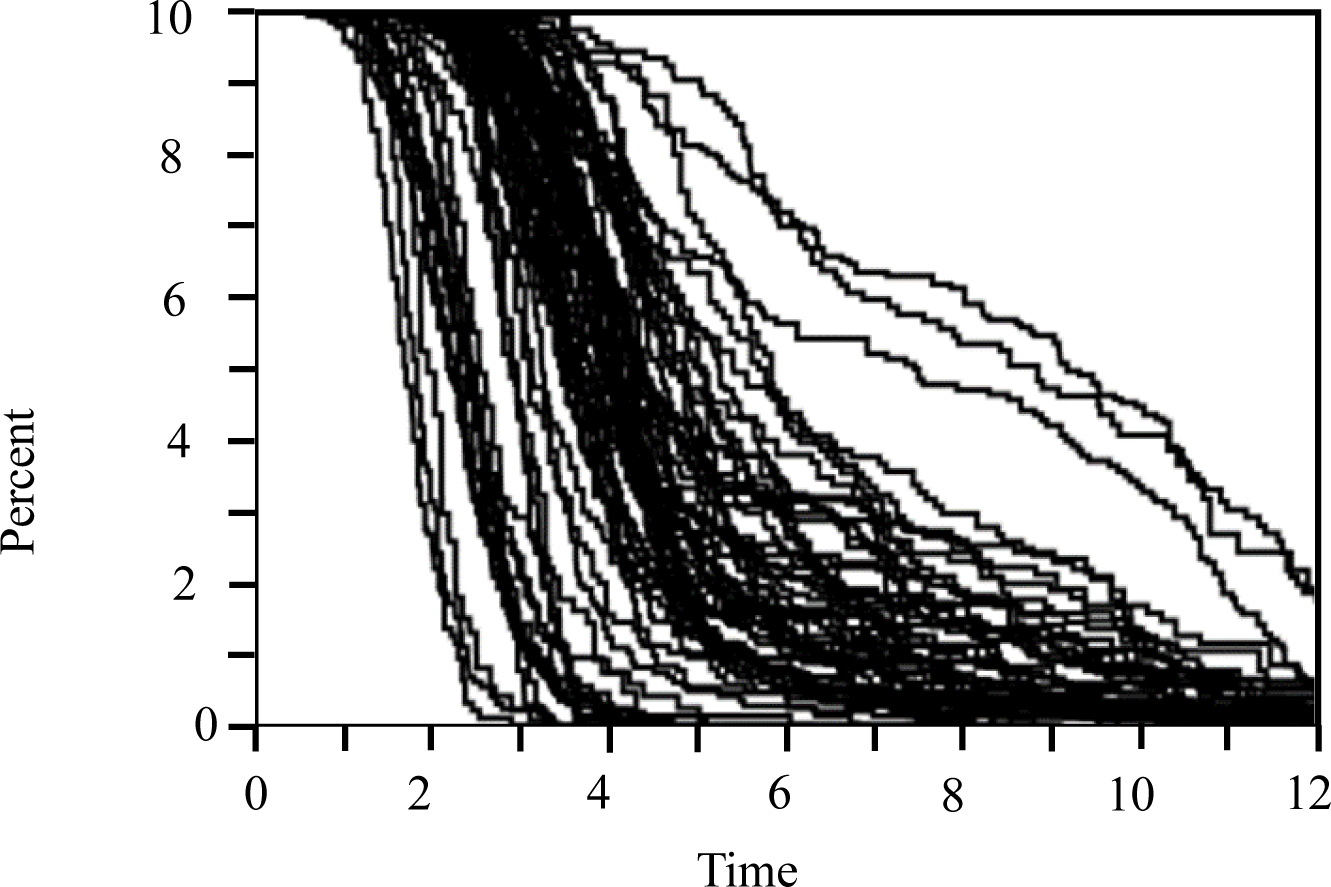
CeMEE panel adult starvation survival curves. Survival curves of 72 CeMEE panel lines. Each curve represents one CeMEE line with technical replicates ≥ 2; replicate assays initiated on different days. At time 0, animals are early adults and within 1 hour of placement into the microfluidic starvation environment. Analysis of Kaplan-Meier lifespan curves with log rank test for statistical differences was performed using JMP Statistical Software (SAS Institute Inc., JMP Pro 13, 2017)

In this system, late L4/very early adults are removed from a plate-based environment with abundant food and placed directly into a microfluidic environment in the absence of food. The typical response of *C. elegans* individuals when food becomes scarce is to seek out a new location where food is present (Shtonda & Avery, 2006; Kang & Avery, 2009). In this experimental setup individuals can attempt to flee as normal but ultimately are retained within the pillared arena until death. The measurements presented here capture the variety of starvation responses in our CeMEE panel lines. The survivorship curves presented in Figure 4 appear to be dividable into two categories based on the curve shapes. The first are curves with a sigmoidal shape. In this group, individuals tend to die off quite rapidly and these comprise lines with lower mean and median survival lengths (Supplemental Table 1). The second group are curves that have a more consistently gradual decrease in shape. The individuals in this group die off slowly and make up the lines with higher mean and median survival times. This is suggestive of two different sources of mortality in response to starvation and is consistent with previous observations of adult *C. elegans* where, when confronted with a loss of food, some hermaphrodites respond by sacrificing themselves to matriphagy and others enter a state of reproductive diapause (Angelo & Van Gilst, 2009; Seidel & Kimble, 2011; Burnaevskiy et al., 2018).

### Genetic Variation and Median Survival

To identify genetic variation contributing to differences in the time-until-death phenotype under microfluidic starvation conditions, we used a GWA mapping approach. For this approach we treated the time to death of each individual as a quantitative trait (n = 7855). Plink and rrBLUP identified a similar set of SNPs as significant contributors to this phenotype but with variance in the absolute levels of significance across the genome (Supplemental Figure 1, Supplemental Table 2). Because of the similarity, only the rrBLUP results will be presented here (Figure 5, Table 1).

**Figure 5:**
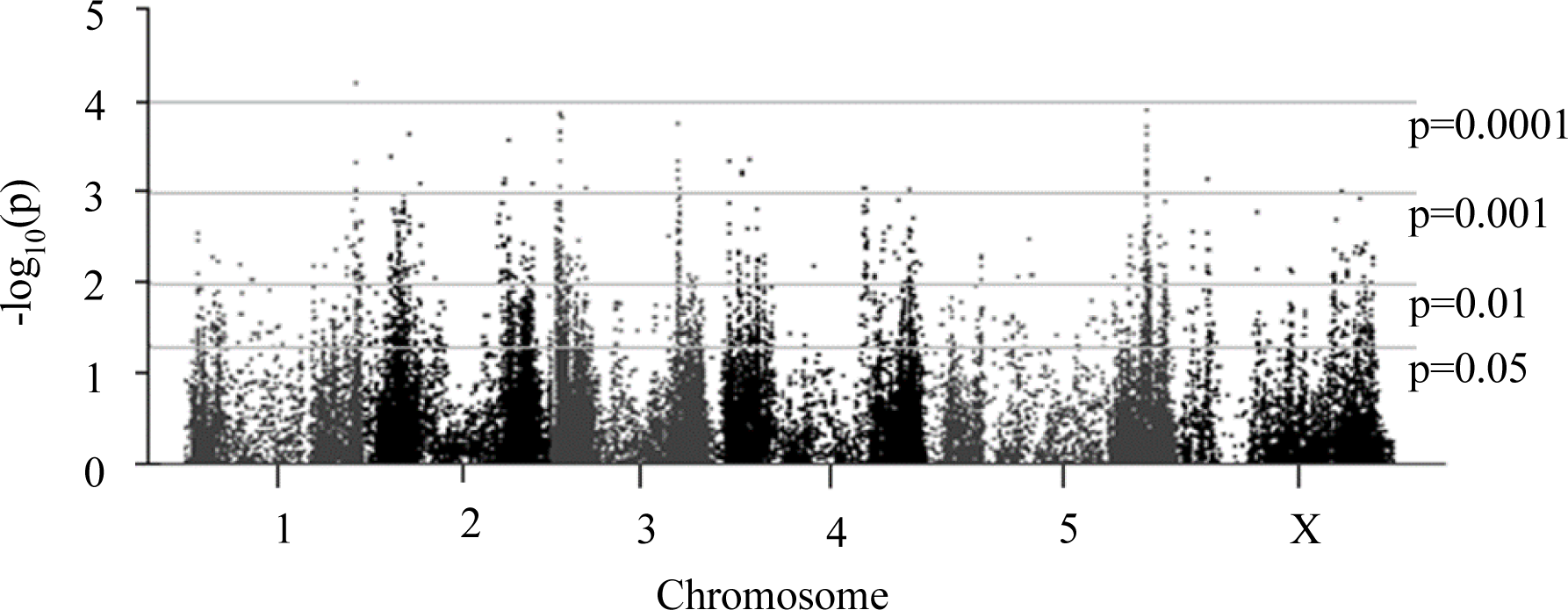
Genome-Wide Association Manhattan plot (rrBLUP). Distribution of variants identified through genome-wide mapping with rrBLUP (p-values corrected for kinship, details in methods).

**TABLE 1.** SNPs identified through GWA mapping as significant contributors to median survival length in the microfluidic starvation environment. SNPs with -log_10_(p) above 3.5 were selected. The cutoff of 3.5 was chosen because it was sufficient to capture the majority of significant SNPs with R^2^ values above 0.10. In a few instances there were multiple adjacent SNPs of equal significance, in these cases the SNP located in the middle was chosen as a representative SNP and the size of the associated region is listed in the Region column. See results section for detailed information on the Chromosome V region. In two instances there were two adjacent SNPs with equal significance values, for these there is an arrow indicating the direction of the region associated with the chosen representative SNP. Associated genes were identified by searching within the specified region or within 1 KB on either side for single SNPs. After examination of the per line median survival times plotted by genotype (Figure 6), the four SNPs in bold have clear directionality and were considered the loci of most significance (also see Supplemental Figure 2). WSXXX denotes reference genome version.

| Chromosome | SNP (WS245) | SNP (WS269) | -log<sub>10</sub>(p) | R<sup>2</sup> | Genes | Region |
| --- | --- | --- | --- | --- | --- | --- |
| I | 14359698 |  | 4.23 | 0.11 | *Y105E8A.3* | Single |
| II | 3704344 |  | 3.66 | 0.05 | *srx-107* | Single |
| **II** | **11986204** | **11986223** | **3.60** | **0.16** | ***ash-2*** | **Single** |
| III | 922537 |  | 3.90 | 0.12 | *pef-1* | Single |
| III | 924366 |  | 3.88 | 0.11 | *pef-1* | Single |
| III | 974499 |  | 3.60 | 0.10 | *R06B10.2* | 108 BP |
| III | 998200 |  | 3.71 | 0.11 | *trp-2, R06B10.7* | 8.2 KB |
| III | 1010961 |  | 3.68 | 0.09 | *Y34F4.2, Y3F4.6* | ←1.5 KB |
| **III** | **1107965** | **1107975** | **3.85** | **0.18** | ***exc-6*** | **Single** |
| **III** | **10768218** | **10768084** | **3.78** | **0.18** | ***dpy-28*** | **2 KB→** |
| V | 18266401 |  | 3.67 | 0.11 | *R10E8.6, fbxa-213* | 4 KB/26 KB |
| **V** | **18271385** | **18271413** | **3.94** | **0.11** | ***Y37H2C.4*** | **Single/26 KB** |
| V | 18277648 |  | 3.75 | 0.12 | *Y37H2C.1, Y37H2C.5* | 2.7 KB/26 KB |
| V | 18293029 |  | 3.54 | 0.12 | *Y51A2A.12* | Single/26 KB |
| V | 18312716 |  | 3.75 | 0.06 | *fbxa-117* | Single |

Potential causative genetic variation underlying each QTL was identified by first extracting all SNPs from each dataset where -log_10_(p) ≥ 3.50, sufficient to select the majority of SNPs with R^2^ > 0.10 (SNPs below this value do not show a clear direction of influence on median survival time, see Supplemental Figure 2). This resulted in a total of 40 focal SNPs, however after eliminating adjacent and redundant SNPs (as described in Table 1), this number was further collapsed to a total of 15 loci. The direction and size of the effect of each locus was determined by calculating the difference in median survival time per allelic class (Figure 6). Eight SNPs identified in our GWA mapping were either heavily influenced by a singleton “outlier” line or generated a genotype/phenotype correlation that did not indicate a clear directionality of effect, and so are not given further consideration here (Supplemental Figure 2). Four of the SNPs on chromosome V are in close proximity and largely redundant (Supplemental Table 3), as discussed in further detail below.

**Figure 6:**
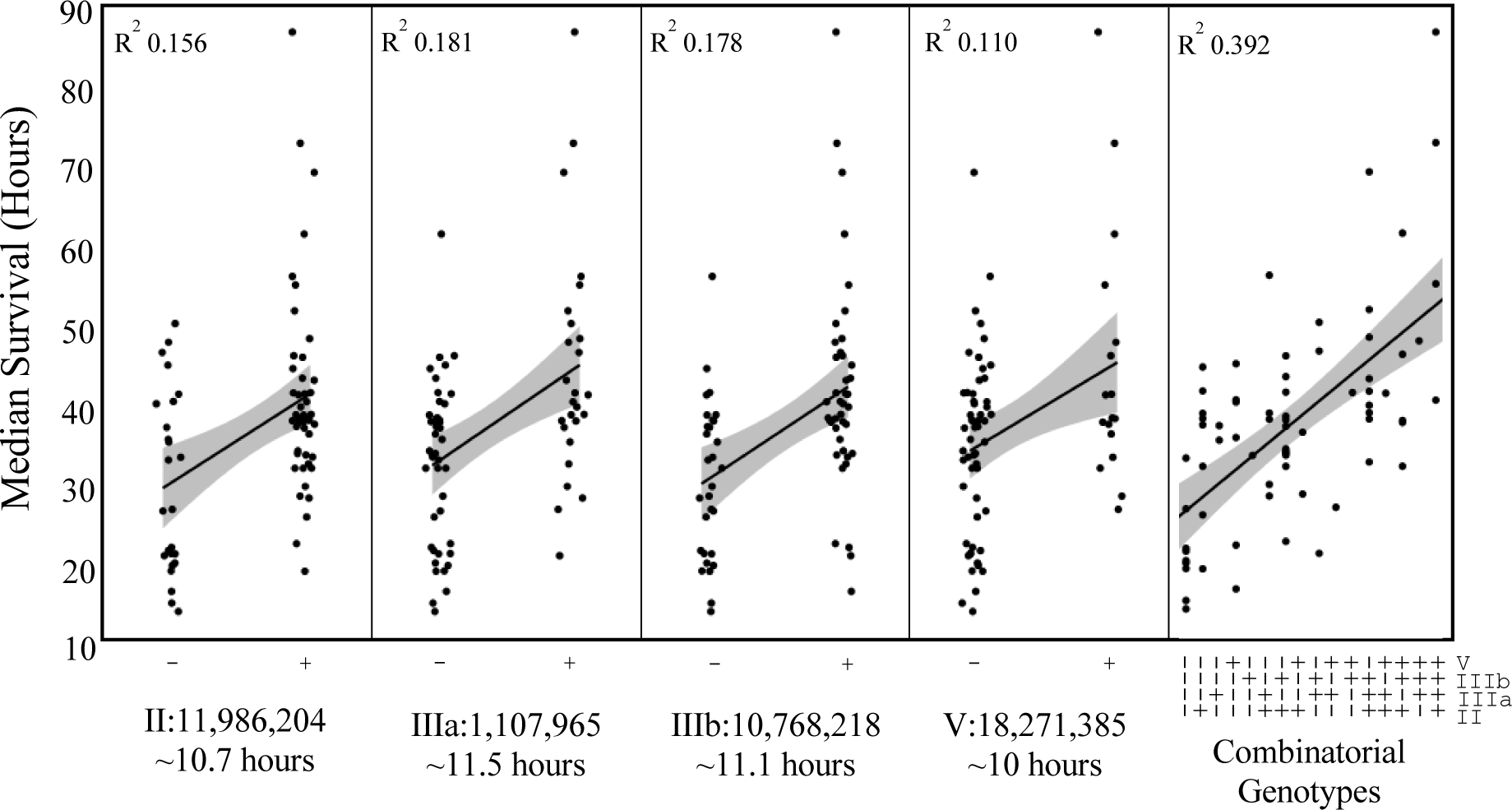
Effect sizes of SNPs contributing to differences in median lifespan under microfluidic starvation conditions. Genotypes denoted as ‘+’ are the N2 reference allele. All alleles are biallelic. Fit line is linear regression (shaded area = 95% confidence interval).

The most suggestive locus is on the left arm of chromosome III (BP 1,107,965). The alternate genotype is present in 46 of the 72 CeMEE lines and represents an average drop in median survival time of ∼11.5 hours. This locus accounts for 18.1% of the observed among-line variation and is a T → A substitution within an intron of the gene *exc-6*.

A locus of nearly equally large effect was identified on the right arm of chromosome III at position 10,768,218. This locus accounts for 17.8% of variation and reflects an average decrease in median survival time of ∼11.1 hours. Thirty of the CeMEE lines have the alternate genotype at this locus, a T → C substitution in an intron of the gene *dpy-28*. In our dataset this locus partitioned equally with a SNP at III: 10,770,501 so these loci are more likely to be representative of an effect coming from the ∼2 KB region spanning from one SNP to the other rather than either individual variant (Table 1).

A third locus of interest is on the right arm of chromosome II at base pair position 11,986,204. This locus represents a ∼10.7 hour decrease in median survival time. Twenty-five of our tested lines have the alternate genotype, a G → A substitution in an intron of the gene *ash-2*. This locus accounts for 15.6% of the variation in median survival time between lines.

A region on the right arm of chromosome V at position 18,271,385 is the final locus of interest. It accounts for 11% of variation and reflects an average decrease in median survival of ∼10 hours. Fifty-five of our tested lines have the alternate genotype, a known missense mutation in the gene *Y37H2C.4*. Moreover, this locus appears to represent the focal point of a ∼26 KB region where multiple genetic variants affect median survival across the CeMEE lines. Three other loci identified as significant in the GWA mapping surround this SNP and have only slightly different membership across the CeMEE panel lines (n = 57, 56, and 54; Table 1, Supplemental Table 3). Due to the difference in membership they have somewhat lower p-values and were initially 5 examined as separate contributory sources. However, on closer examination it became apparent that the only time a significant increase in median survival was seen was when the first three variants collectively match the reference genotype. The fourth variant, V:18,293,029, does not appear to contribute any additional effect (see Supplemental Figure 3 for side-by-side comparison of 3- and 4-locus combinatorial genotype effect sizes) and so it is not included in further analysis. Of our 72 lines, 53 have the alternate genotype at all three loci and 5 have an intermediate combination. There does not appear to be any significant difference between these lines (Supplemental Figure 3). However, the remaining 14 lines have the reference genotype at all 3 loci and this group displays the ∼10 hour increase in median survival time. This is not to say that any individual SNP exerts a greater influence or that a specific combination is necessarily meaningful, more intermediate data points are needed, this is only to suggest that it is variation in this region as a whole influencing the survival phenotype.

Lastly, when the above four individual loci are examined collectively as combinatorial genotypes, 39.2% of the variation in median survival is accounted for (Figure 6).

### Backcross with Selection

Because of the crossing strategy used to create the CeMEE mapping panel, each line represents a mosaic of variants segregating in natural populations. However, because of the multiple generations of inbreeding by selfing after the initial crossing, the variants are largely fixed within lines, with each line largely representing a static mosaic of fixed variants spread throughout the genomes (Figure 1). With this in mind we chose to use a backcross with selection strategy to independently isolate the variants contributing to starvation response. We chose one line which displayed the slow-dying/long-lived phenotype and another which displayed the fast-dying/short-lived phenotype (Figure 4, Supplemental Table 1) to be the founding parents. We then continually selected for the long-lived phenotype while repeatedly crossing into the short-lived genome as described above.

The near-isogenic lines derived from the backcross with selection population segregated into two phenotypic groups matching the respective parent phenotype in a ratio of 11:6 long to short, with an average difference of 10 hours in median lifespan (Figure 7). Comparison of the genomic sequences from 8 NILs (4 long-lived and 4 short-lived) plus the parent lines shows a region on the right arm of chromosome V, starting around base 18,200,000 and continuing to the end of the chromosome containing variants common to only the long-lived NILs and long-lived parent (Figure 8). A comparison across the two groups (short-vs long-lived NILs) gives a Cohen’s D of 0.81, suggesting this is a locus of large effect and supporting the identification of this locus as a significant contributor to median survival in the GWA mapping approach.

**Figure 7:**
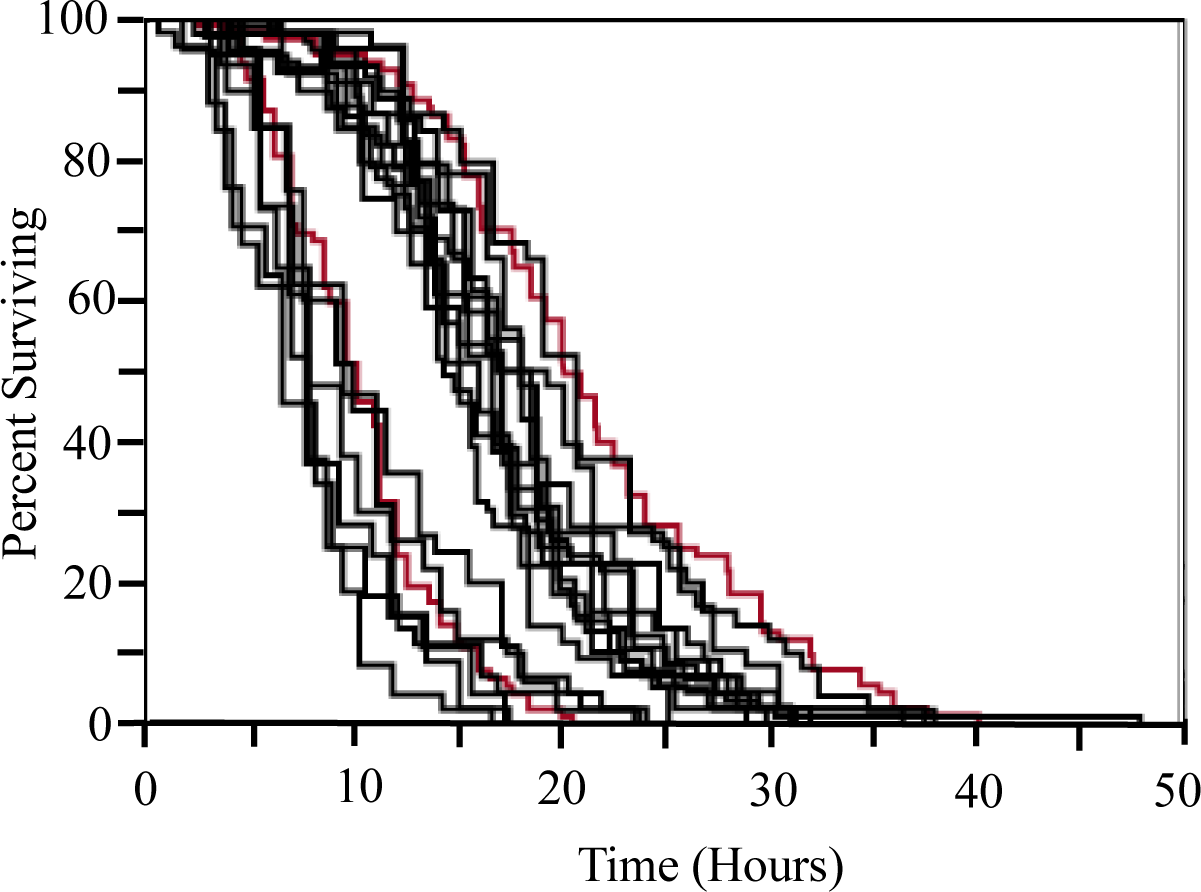
Survival curves of adults from the near-isogenic lines in the microfluidic starvation environment. Adult hermaphrodite survival was measured in the same manner as previously described. The near-isogenic lines segregated into a ratio of 11:6 (long:short) with an average mean difference of ∼10 hours. Black lines are near-isogenic lines, red lines are the CeMEE parental lines (A6140L110 and GA450L40).

**Figure 8:**
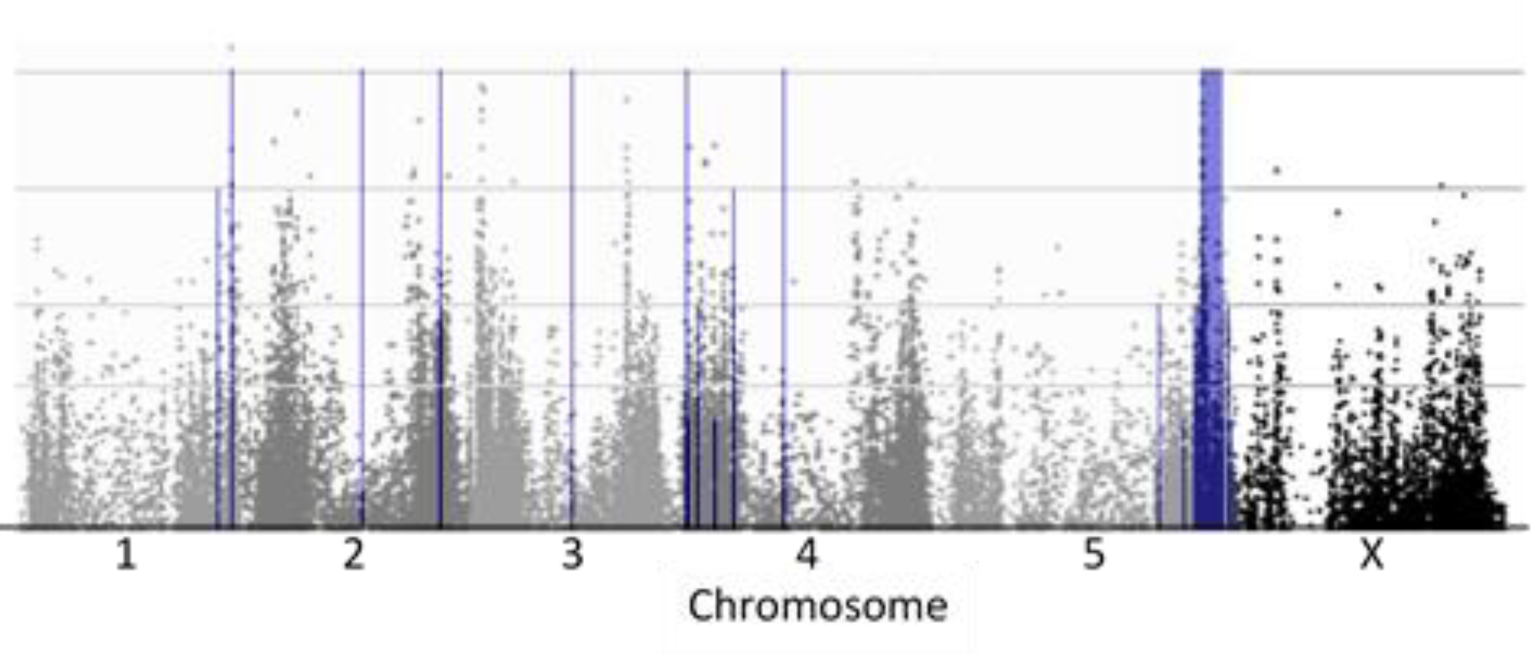
Chromosomal regions from long-lived parent retained by long-lived NILs. Dark blue regions represent the portions of each chromosome shared between the long-lived NILs and the long-lived parent exclusively. The height of the dark blue bar reflects the number of long-lived NILs (1-4) matching the parent sequence. The block on Chromosome V starts around base 18,200,000 and extends to the end of the chromosome. An overlay of the GWA Manhattan plot (Figure 5) with the long-lived parent chromosomal regions retained in the NILs shows the same regions of significance.

Within this region there is fine-scale variation in which the long-lived NILs possess variants matching only the short-lived parent and/or novel mutations in between small blocks of the long-lived genotype (Supplemental Figure 4). As such, recombination has occurred with retention of the presumably necessary variants from the long-lived parent.

## Discussion

Organisms respond in a multitude of ways when faced with resource limitation. It has been well-established in *C. elegans* that dietary restriction and food deprivation can significantly extend overall lifespan (see Uno & Nishida, 2016 for a comprehensive review). In early larval stages (L3 and prior) individuals can arrest or adopt alternate development strategies to cope, allowing them to stay in these states for an extended amount of time until food becomes sufficient and development resumes as normal (Klass & Hirsh, 1976; Johnson, Mitchell, Kline, Kemal, & Foy, 1984; Fielenbach & Antebi, 2008; Zhou, Pincus, & Slack, 2011; Baugh, 2013; Schindler, Baugh, & Sherwood, 2014; Roux, Langhans, Huynh, & Kenyon, 2016). When food is reduced or lost during later days of adulthood (day 2 and after) individuals also typically have an increase in lifespan over their well-fed counterparts (Kaberlein et al., 2006). However, the picture is not quite as clear during the last larval stage (L4) and very early adulthood. This is the time during which *C. elegans* directs much of its energy into reproductive processes under normal circumstances and it has been shown that loss of food during this period can lead to very different outcomes (Angelo & Van Gilst, 2009; Seidel & Kimble, 2011; Burnaevskiy et al., 2018). Based on the expression patterns of genes involved in vulval development, the L4 larval stage can be roughly divided into three substages; early (hours 0-4), middle (hours 4-8), and late (hours 8-9) (Mok, Sternberg, & Inoue, 2015). While the effect of food removal on development through these substages is not entirely clear, it is evident that when food is removed during late L4 and/or the transition into adulthood, as was done in this study, individuals either undergo facultative vivipary (where eggs are fertilized but not laid, thereby hatching within the body of the parent hermaphrodite, also referred to as ‘bagging’), or enter a state of adult reproductive diapause (Angelo & Van Gilst, 2009; Seidel & Kimble, 2011; Burnaevskiy et al., 2018). The two different shapes of curves seen in Figure 4 likely reflect this – the ‘rapid’ death of the sigmoid shape reflects lines where the majority of individuals bagged and diapause lines are those with the comparatively ‘slow’ deaths of the gradually decreasing slope.

Our GWA mapping identified four loci with clearly additive effects underlying the variation in starvation response. Two of the loci are individual SNPs within introns of the genes *ash-2* and *exc-6*, the other two are representative of larger haplotype blocks: 1) a ∼2 KB region associated with the gene *dpy-28* and 2) a ∼26 KB region encompassing several genes on Chromosome V (Table 1). Because linkage in our tested lines is low, the genetic components underlying the variation seen here are expected to be influencing the trait in a mostly additive fashion. If this is true, then the expectation is that for any line with a genotype containing a combination of two or more advantageous alleles, the increase in lifespan over those without the alleles should equal the sum of the increase for those with the respective single allele:

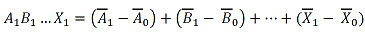

This appears to be the case. Figure 6 shows the best fit line and the median survival time values for all genotype combinations in this dataset. While the overall increase predicted by the best fit line is approximately 30 hours, the observed increase across the mean values for the two extremes is 42.3 hours which nearly matches the strictly additive model prediction (10.7 + 11.5 + 11.1 + 10 = 43.3). However, more information from lines composed of double and triple genotype combinations would be needed to confirm this with certainty.

Consistent with the lifespan curves, animals recovered from the microfluidic devices during the backcross with selection had either bagged or displayed the very poor morphological appearance with reduced intestines and somatic gonads, similar to the description of animals in adult diapause described in Angelo & Van Gilst (2009), when first placed on recovery plates. Moreover, as in the Angelo & Van Gilst (2009) study, this poor morphological appearance gave way to a fully rejuvenated appearance by recovery day two in the individuals who produced offspring. This suggests that we have mapped a naturally segregating polymorphism comprising two distinct responses to reduced nutrient availability: transition to self-consumptive reproduction or enter diapause awaiting a change in the environment.

While our results suggest that individuals can enter a state of adult reproductive diapause in the microfluidic starvation environment, there is one striking difference between the individuals in this study and previous studies in which individuals have been reported to remain in a diapause state for up to at least three weeks, i.e. 504 hours (Burnaevskiy, et al., 2018). In our microfluidic starvation environment, maximum lifespan appears to be closer to ∼140 hours than three weeks. There are several, not mutually exclusive, possibilities for this difference. 1) The previous studies were conducted on unseeded agar plates that typically contain antibiotics to suppress the growth of bacteria whereas our microfluidic setup uses constant perfusion of sterile fluid to create an environment devoid of bacteria and other substances, and so there may be an effect of the plate environment. 2) When on plates, animals are in contact with each other and presumably take chemical cues from one another (Kaplan et al., 2011), which may influence the diapause state, whereas in our microfluidic chip animals are reared separately and constant perfusion means that any chemical forms of communication are washed away. 3) Data on adult reproductive diapause has been collected primarily, if not exclusively, from animals with the N2 genetic background in which case adult diapause as reported may be unique to that background as opposed to the naturally-derived backgrounds used here. 4) Adult reproductive diapause may be a complex trait involving the cohesive (linked) function of multiple loci that have become uncoupled during the creation of the CeMEE panel (Figure 1).

While the backcross with selection was intended to be a mapping approach parallel to GWA, the selection approach actually imposed a slightly different phenotypic criterion than was measured for GWA. In the GWA approach, adult individuals were placed into the starvation environment and monitored without further intervention until all were dead. In the backcross with selection, adult individuals were placed into the starvation environment and monitored until ∼20% of the population remained or 24 hours had passed, whichever came first. Surviving individuals were then removed and the fraction which produced offspring are the genotypes that progressed to the next backcross. Therefore, when survival length was measured for the backcross with selection, individuals had been selected to 1) have an increased length of survival and 2) be able to produce offspring when food was reintroduced after ∼80% of the population had succumbed to starvation. Nevertheless, an overlay of the region retained in the long-lived NILs and the significant chromosome V region from the GWA shows that the regions completely overlap, at least to the fairly limited linkage resolution of the backcross with selection approach (Figure 8). It is likely that total length of survival and ability to produce offspring post-starvation are related but different traits, therefore it is not necessarily surprising that variants from 3 out of the 4 loci significant in the GWA approach were not retained in the long-lived genotypes resulting from the backcross with selection.

The retained region spans ∼2.7 MB, beginning around position V:18,200,000 while the GWA locus has the approximate genomic position of V:18,264,009-18,293,029, a length of ∼26 KB. This is the locus contributing an average 10-hour increase in median lifespan among the CeMEE panel lines and the same average difference is seen between the short- and long-lived NILs. If linkage and epistatic interactions between this locus and other genomic regions are truly diminished (as expected for lines from the CeMEE panel), then the phenotype resulting from introgression of this locus into a different genetic background would be expected to have only the additive effect. Given that the average change in median survival is the same in the NIL lines as it is for lines in the CeMEE panel, then this 1) reaffirms the reduction/loss of linkage disequilibrium in the CeMEE panel and 2) confirms this effect as additive (Wright, 1952; Hill, 1998; Luo, WU, & Kearsey, 2002; Hospital, 2005). Moreover, this chromosome (and more specifically this region) has the only sign of linkage drag. None of the other chromosomes have retained significant portions of the long-lived parent genome (Figure 8). Moreover, it is likely that variants within this locus have a directed effect on post-starvation reproductive ability.

Although backcross with selection as a technique for isolating QTL has long history in the literature, starting from the first proposal by Wright (1952) (and expanded to include *inter se* mating as was done here by Hill in 1998), it has not been extensively used as a primary mapping approach. Here, backcross with selection was not used as the primary mapping approach, but was intended to verify loci previously identified through GWA mapping. Subtle differences in phenotype highlight potential challenges in applying different mapping approaches to the same question, although the overlap that we observe here provides confidence that both the observed effects and mapping locations are valid for this particular locus. A recent similar study used a population selection/RAD-seq approach to quantify genetic variation and analyze starvation resistance in *C. elegans* at the L1 larval stage (Webster et al., 2019). The results show substantial natural variation in 96 wild isolates from the *C. elegans* Natural Diversity Resource (CENDR) panel and identified an associated region on the left arm of chromosome III using GWA mapping. However, in this case the authors validate the genotype/phenotype association by creating a NIL through introgression of the chromosome III region from a starvation-sensitive strain into the genetic background of a starvation-resistant strain, effectively selecting on the basis of genotype rather than phenotype. This differs from our methodological approach of using selection in a backcross introgression scheme. To our knowledge this is the first instance of our technique being applied in this manner to *C. elegans*.

The three loci identified in the GWA mapping that are absent from the backcross results have interesting physiological potential as candidates for influencing overall lifespan through starvation response. The gene *ash-2*, associated with the locus at II:11,986,204, is part of a histone methyltransferase complex known to be involved in regulating adult lifespan through its effects on H3K4 methylation in the *C. elegans* germline (Greer et al., 2010; Greer et al., 2011; Zuryn et al., 2014; Robert et al., 2014). Given that this locus already has a known role in affecting lifespan through reproduction, further investigation of variation in this gene under the context of starvation response is warranted. The gene *exc-6*, associated with the III:1,107,965 locus, is expressed throughout the pharynx, rectal gland cells, and the reproductive system in both larval and adult individuals and localizes to filamentous actin (Shaye & Greenwald, 2016; Hegsted, Wright, Votra, & Pruyne, 2016). However, it does not appear to have any previously reported role in affecting lifespan or starvation response, but *exc-6* mutants do exhibit defects in ovulation, suggesting this locus does have a significant role in reproduction and may influence survival in a starvation environment by restricting ovulation (Hegsted, Wright, Votra, & Pruyne, 2016). It would be interesting to examine the relationship between variation in *exc-6* and the bagging/ARD decision as presumably reduced ovulation would push energy investment in the direction of somatic maintenance. The third locus, at III:10,768,218, is associated with the gene *dpy-28*. This gene is involved in meiotic sister chromatid segregation, mitotic sister chromatid segregation, negative regulation of reciprocal meiotic recombination, and localizes to the intracellular organelle (Hernandez, et al., 2018). Interestingly, mutants of *dpy-28* have been reported to be unable to enter the dauer arrest state via an effect on the transcription factor DAF-16/FOXO, a widely recognized stress response pathway with dramatic effects on lifespan and starvation response in *C. elegans* (Weinkove, Halstead, Gems, & Divecha, 2006; Dumas et al., 2013; Uno & Nishida, 2016). The genetic basis for this is suggested to be mediated by a role of *dpy-28* in dosage compensation, such that *dpy-28* mutants may have elevated expression of X-linked genes that typically promote dauer bypass. This raises the possibility that variants at this locus may promote the bagging phenotype by making it difficult or preventing individuals from entering into an arrested state in response to starvation cues. Of course, any of these putative effects would need to be confirmed via genetic transformation, and it is formally possible that some of these SNPs could influence the regulation of genes that are not directly adjacent to them (Pastinen, 2010; 1000 Genomes Project Consortium, 2012).

Overall, then, we identified four loci with significant effects contributing to variation in survival time in a microfluidic starvation environment through a GWA mapping approach. One of the four loci, found on the right arm of chromosome V, was confirmed in a parallel backcross with selection approach. The absence of selection on any of the other three loci suggest that they were either lost by chance during the backcross with selection or have different effect sizes under the slightly different criteria – the ability to reproduce post-extended starvation (Hill, 1998; Hospital, 2001; Hospital, 2005). Taken together this suggests that the absolute adult lifespan under starvation conditions may be partially independent of the ability to recover and reproduce after surviving extended starvation and the locus identified in both approaches may act in a pleiotropic fashion to influence both traits. When faced with a loss of food during the period of time during which *C. elegans* hermaphrodites normally balance a physiological investment of energy into maturation of the germline with investment into somatic maintenance, individuals have to make a choice of where to partition their energetic resources. Some individuals are genetically predisposed to sacrifice their soma and undergo facultative matricide while others can maintain their soma and effectively put germline maturation on pause, suggesting that the natural variation segregating in populations of *C. elegans* may be evolving two different starvation response strategies.

## Acknowledgments

A special thanks to Luke M. Noble for discussion, ideas, collaboration, and support. We thank John Willis for preparing the sequencing library used in this research and Henrique Teótonio for providing the CeMEE strains. We would also like to thank John Conery, Bill Cresko, Mike Harms, and John Postlethwait for their support and guidance. Justin Robles assisted in establishing a method for handling and processing this data set. This work was supported by the National Institutes of Health (T32 training grant GM007413 to HA, R01 AG049396 and R01 GM102511 to PCP).

